# Handling of intracellular K^+^ determines voltage dependence of plasmalemmal monoamine transporter function

**DOI:** 10.1101/2021.03.11.434931

**Authors:** Shreyas Bhat, Marco Niello, Klaus Schicker, Christian Pifl, Harald H. Sitte, Michael Freissmuth, Walter Sandtner

**Author notes:** S. Bhat and M. Niello contributed equally to this paper. Correspondence: Walter Sandtner.

## Abstract

The concentrative power of the transporters for dopamine (DAT), norepinephrine (NET) and serotonin (SERT) is thought to be fueled by the transmembrane Na^+^ gradient, but it is conceivable that they can also tap other energy sources, e.g. membrane voltage and/or the transmembrane K^+^ gradient. We address this by recording uptake of endogenous substrates or the fluorescent substrate APP^+^ ((4-(4-dimethylamino)phenyl-1-methylpyridinium) under voltage control in cells expressing DAT, NET or SERT. We show that DAT and NET differ from SERT in intracellular handling of K^+^. In DAT and NET, substrate uptake was voltage-dependent due to the transient nature of intracellular K^+^ binding, which precluded K^+^ antiport. SERT, however, antiports K^+^ and achieves voltage-independent transport. Thus, there is a trade-off between maintaining constant uptake and harvesting membrane potential for concentrative power, which we conclude to occur due to subtle differences in the kinetics of co-substrate ion binding in closely related transporters.

## Introduction

The dopamine transporter (DAT), the norepinephrine transporter (NET) and the serotonin transporter (SERT) are members of the solute carrier 6 family (*SLC6*). DAT, NET and SERT mediate reuptake of released dopamine, norepinephrine and serotonin, respectively (**Kristensen et al., 2011**). By this action, they terminate monoaminergic signaling and – in concert with the vesicular monoamine transporters – replenish vesicular stores. These transporters are secondary active in nature; they utilize the free energy contained in the transmembrane Na^+^ gradient (established by the Na^+^ /K^+^ pump) to drive concentrative monoamine uptake into cells in which they are expressed (**Burtscher et al., 2019**). *SLC6* transporters have been postulated to operate via the alternate access mechanism (**Jardetzky, 1966**): they undergo a closed loop of partial reactions constituting a complete transport cycle (**Rudnick & Sandtner, 2019**). These partial reactions require conformational rearrangements and binding/unbinding of substrate and co-substrate ions. It is gratifying to note that crystal structures obtained from the prokaryotic homolog LeuT, drosophila DAT and human SERT itself support the general concept of alternate access (**Yamashita et al., 2005; Penmatsa et al., 2013; Coleman et al., 2016**). These crystal structures also reveal SERT and DAT to be closely related. This is evident from the root mean square deviation, which differs by only approximately 1 Å between the outward facing structure of human SERT and drosophila DAT. DAT, NET and SERT also share a rich and partially overlapping pharmacology (**Sitte and Freissmuth, 2015**): there are many drugs that inhibit transporter function by either blocking or inducing reverse transport. These mechanisms account for therapeutics used for the treatment of neuropsychiatric disorders (major depression, general anxiety disorder and attention-deficit hyperactivity disorder) and for many illicit drugs that are psychoactive and abused (**Hasenhuetl et al., 2019; Niello et al., 2020**).

Despite the similarity in structure and function, the three transporters differ in many more aspects than just ligand recognition: the transport stoichiometry of SERT and DAT/NET is considered to be electroneutral and electrogenic, respectively. It has long been known that SERT antiports K_in_^+^, i.e., intracellular K^+^ (**Rudnick & Nelson, 1978**); for DAT and NET, K_in_^+^ is thought to be immaterial (**Gu et al., 1994; Gu et al., 1996; Sonders et al., 1997; Erreger et al., 2008**). If true, only SERT can utilize the chemical potential of the cellular K^+^ gradient to establish and maintain a substrate gradient. It has therefore remained enigmatic, why closely related transporters can differ so fundamentally in their stoichiometry and their kinetic decision points. The effects of K_in_^+^, intracellular Na^+^ (Na_in_^+^) and membrane voltage on the transport cycle of SERT have been recently analyzed in great detail (**Hasenhuetl et al., 2016**). However, much less is known on how these factors impinge on the transport cycle of DAT and NET (**Galli et al., 1998**, **Sonders et al., 1997**; **Hoffman et al., 1999**; **Prasad and Amara, 2001**; **Erreger et al., 2008; Li et al., 2015**). In this study, we investigated the role of intracellular cations and voltage on substrate transport through DAT, NET and SERT. To this end, we simultaneously recorded substrate induced currents and uptake of the fluorescent substrate APP^+^ (4-(4-dimethylamino)phenyl-1-methylpyridinium) into single HEK293 cells expressing DAT, NET and SERT under voltage control. These measurements were conducted in the whole cell patch clamp configuration, which allowed for control of the intra- and extracellular ion composition via the electrode and bath solution, respectively. Our analysis revealed that K_in_^+^ did bind to the inward facing state of DAT and NET but, in contrast to SERT, K_in_^+^ was released prior the return step from the substrate free inward-to the substrate free outward-facing conformations. We also found that substrate uptake by DAT and NET, unlike SERT, was voltage-dependent under physiological ionic gradients. Moreover, the absence of K_in_^+^ had no appreciable effect on the transport rate of DAT and NET. The transient nature of K_in_^+^ binding was incorporated into a refined kinetic model, which highlighted the differences between SERT and DAT/NET. Notably, this model allows for a unifying description, which attributes all existing functional differences between DAT, NET and SERT to the difference in the handling of K_in_^+^.

## Results

### Single cell uptake of APP^+^

We combined advantages of transporter-targeted radiotracer assays and electrophysiology by setting up a system wherein APP^+^ (*Fig. 1A, left*) mediated uptake through a single DAT, NET or SERT-expressing HEK293 cell was measured under voltage control (*Fig. 1A, center*). *Fig. 1A, right* is a theoretical representative of the two channel recordings; the orange trace represents time dependent-increase in fluorescence (i.e., APP^+^ accumulation intracellularly), while the red trace represents APP^+^-induced currents. *Fig. 1B* are actual representative traces of the two-channel recordings obtained from control (untransfected) HEK293 cells or from HEK293 cells expressing DAT, NET or SERT voltage-clamped at −60 mV and exposed to 100 μM APP^+^. In control HEK293 cells (left hand set of traces in Fig. 1B), there was a transient sharp increase and decline in fluorescence as APP^+^ is washed-in and washed-out, respectively. There wasn’t any concomitant change in current. The rapid rise in fluorescence most likely represents reversible binding of APP^+^ to the plasma membrane. It was also seen in HEK293 cells expressing transporters. In DAT-expressing cells, APP^+^ induced an inwardly directed current comprised of both, a peak and a steady state component, indicating that APP^+^ is a DAT-substrate. Transport of APP^+^ into DAT-expressing cells led to a slow rise in fluorescence. This increase was linear with time and terminated upon removal of APP^+^ : fluorescence relaxed to a new baseline, which indicated intracellular trapping of APP^+^. In contrast, APP^+^ induced only inwardly directed peak currents (but no steady state currents) in NET or SERT-expressing cells. Nonetheless, SERT or NET-expressing cells show a linear increase in APP^+^ accumulation over time in these cells, indicative of APP^+^ transport.

**Fig. 1.**
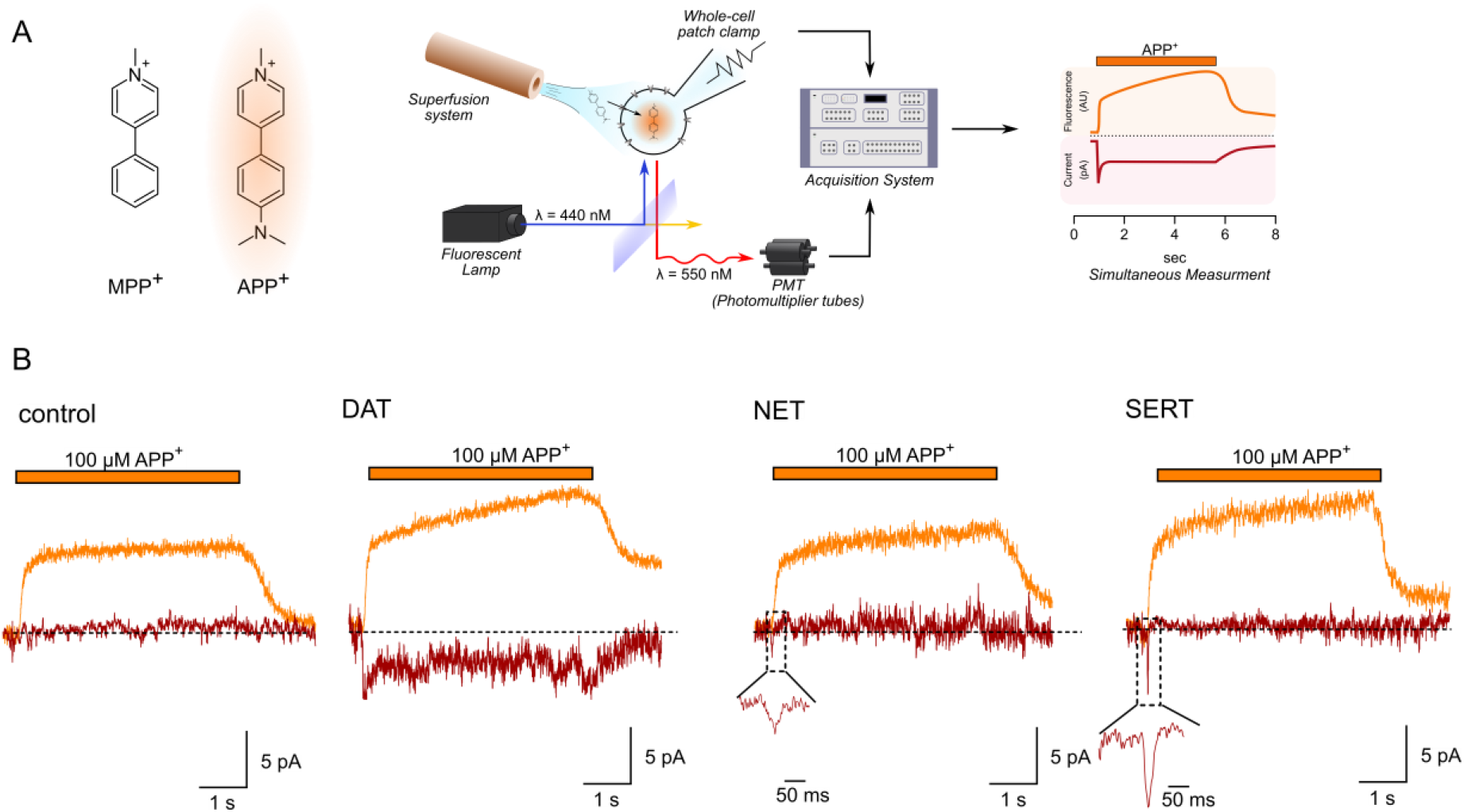
(*A*) *left*, APP^+^ is a fluorescent analogue of MPP^+^. Excitation and emission spectra of APP^+^ occurs at wavelengths ranging from 410-450 nm and 510-550 nm, respectively, which depend on solvent properties. *Center*, Schematics of the setup used in this study. The setup included: 1) a dichroic glass that specifically reflects light of 440 nm but allows transmission of those with wavelength of 550 nm, 2) an inverted microscope with a 100x objective (for single cell fluorescence recordings), 3) an electrode and patch clamp amplifier that allows for voltage control, 4) a photomultiplier tube (PMT) that converts emitted fluorescence into electrical signal and 5) and an acquisition system that allows for filtering, digitizing and two-channel recordings of APP^+^ induced fluorescence and currents. *Right*, A theoretical trace for a two-channel recording that displays simultaneous APP^+^-induced currents (red trace) and fluorescence (orange trace) in real time. (*B*) Representative traces of the APP^+^-induced currents and fluorescence in empty HEK293 cells or in HEK293 cells expressing DAT, NET and SERT patched under normal physiological ionic conditions. In all traces, the rapid rise and decline in fluorescence on applying and washing off APP^+^, respectively, is probably indicative of APP^+^ adherence to the plasma membrane and non-specific in nature. All three transporters show linear increase in fluorescence as a function of time, indicative of APP^+^ accumulation in cells expressing DAT, SERT and NET. APP^+^ induces robust DAT-mediated currents that comprise both peak and steady state currents. Only peak currents (represented as magnified inset) were seen on APP^+^ application to cells expressing NET and SERT. AU – arbitrary units.

### Concentration dependence of APP^+^-induced currents and fluorescence

The slope of the linear increase in fluorescence has the dimension of a rate (i.e. fluorescence*s^−1^) and is hence a suitable readout for the uptake rate of APP^+^. We therefore determined the concentrations required for achieving half-maximal uptake rates and measured the concomitant currents at −60 mV; original representative traces from single cells expressing DAT, NET and SERT are shown in panels *A&D, B&E*, and *C&F*, respectively of *Fig.2*. In DAT, the APP^+^-induced currents increase over the same concentration range as the rise in the rate of fluorescence. Accordingly, the K_M_-values, which were estimated from fitting the data to a hyperbola (K_M_ = 27.7 ± 7.1 μM and 21.5 ± 10 μM, respectively), were indistinguishable within experimental error (*Fig. 2J*).Dopamine induced transporter mediated currents (*Fig. 2G*) with a K_M_ of 4.4 ± 1.4 μM. Thus, when compared to dopamine, APP^+^ is a low-affinity substrate of DAT (*Fig.2J*). In SERT, APP^+^ did not elicit any appreciable steady state currents (even at the highest concentration tested, i.e., 600 μM), but a robust concentration-dependent increase in peak current amplitudes (*Fig. 2F*).APP^+^, nonetheless, accumulated intracellularly in SERT-expressing cells (*Fig. 2C*) indicating that APP^+^ was a substrate of SERT, which was translocated inefficiently. The K_M_ for uptake of APP^+^ (32.0 ±13 μM) was about an order of magnitude higher than the K_M_ of 5-HT (3.6 ± 1.4 μM) estimated from 5HT-induced steady state currents (*Fig. 2L*). This indicates that APP^+^ uptake is also a low-affinity substrate of SERT. In cells expressing NET, APP^+^ accumulated in a concentration-dependent manner (*Fig. 2B*). Electrophysiological resolution of NET associated currents revealed that neither norepinephrine (*Fig. 2H*) nor APP^+^ (*Fig. 2E*) elicited any steady state currents in NET on rapid application. In addition, APP^+^-induced peak currents were considerably smaller than peak currents elicited by norepinephrine (cf. *Fig. 2E and 2H*). These observations can be rationalized by the following hypothetical explanation: (i) NET cycles at rates considerably slower than SERT and DAT (further explored below), thus explaining the lack of substrate-induced steady state currents (ii) APP^+^ is a low affinity substrate for NET (K_M_ = 37.3 ± 17 μM, *Fig. 2K*), but it nevertheless accumulates in NET-expressing cells at appreciable levels, which allow for fluorescence detection.

**Fig. 2.**
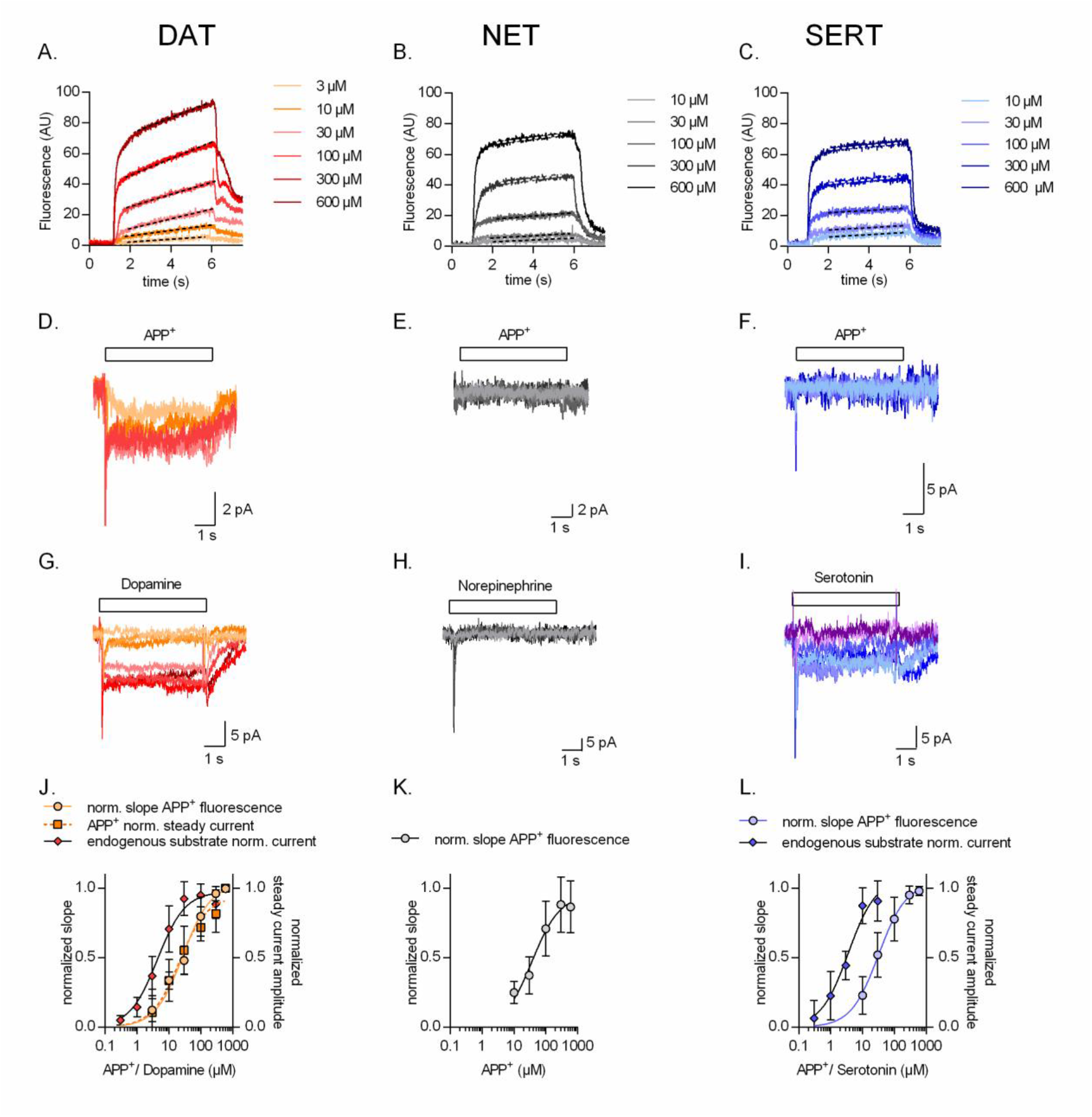
Representative single cell traces of concentration dependent increase in APP^+^-induced fluorescence (*A-C*) and concomitant currents (*D-F*) in DAT, NET and SERT, respectively. Representative single cell traces of currents induced by increasing concentrations of dopamine on DAT (*G*), norepinephrine on NET (*H*) and serotonin on SERT (*I*). (*J*) Normalized concentration response of APP^+^-induced fluorescence (K_M_ = 27.7 ± 7.1 μM (n = 11)), APP^+^-induced steady state currents (K_M_ = 21.5 ± 10 μM (n = 6)) and dopamine-induced steady state currents (K_M_ = 4.4 ± 1.4 μM (n = 6) in DAT-expressing HEK293 cells. (*K*) Normalized concentration response of APP^+^-induced fluorescence (K_M_ = 37.3 ± 17 μM (n = 9)) in HEK293 cells expressing NET. Neither norepinephrine nor APP^+^ induce any steady state currents in NET (even at the highest concentration tested, i.e., 600 μM). (*L*) Normalized concentration response of APP^+^-induced fluorescence (K_M_ = 32.0 ± 13 μM (n = 6)) and serotonin-induced steady state currents (K_M_ = 3.6 ± 1.4 μM (n = 9)) in SERT-expressing HEK293 cells. APP^+^ did not induce any steady state currents in SERT (even at the highest concentration tested, i.e., 600 μM). All experiments were performed under physiological ionic conditions. All fluorescence and current amplitudes were normalized to those obtained at 600 μM (which was set to 1) and the data points were fitted with a rectangular hyperbola. All datasets are represented as means +S.D. AU – arbitrary units, norm. – normalized.

### Voltage dependence of APP^+^-induced currents and uptake

The data summarized in *Fig. 2* indicate that APP^+^ is a substrate, which is taken up with comparable K_M_ by DAT, NET and SERT. Accordingly, we applied APP^+^ for 15 s at a concentration corresponding to the K_M_ range (30 μM) to examine, how changes in voltage (−90 mV to + 30 mV) affect APP^+^ uptake by DAT, NET or SERT-expressing cells in the presence of physiological ionic gradients. It is evident from *Fig. 3A* (representative single cell trace for DAT) and *3B* (representative single cell trace for NET) that DAT and NET show the highest rate of APP^+^ uptake at – 90 mV. The rate of uptake progressively decreased at more positive voltage. In contrast, the change in voltage only had a very modest effect on APP^+^ uptake by SERT (see *Fig. 3C* for a representative single cell trace for SERT). The slopes acquired over the voltage range were normalized to that observed at – 90 mV for each cell and plotted as a function of voltage (circle symbol in Fig. 3 *J, K* and *L* for DAT, NET and SERT, respectively). The plots indicate that uptake of APP^+^ at + 30 mV through DAT, NET and SERT are ~25%, 35% and 80%, respectively, of that at – 90 mV. In control cells, changes in voltage did not affect background APP^+^ binding (data not shown). We also assessed the impact of voltage on steady state currents induced by APP^+^ (representative traces in Fig. *3D-F*) and of the cognate substrates (representative traces in *Fig. 3G-I*). Only, APP^+^ and dopamine evoked transport-associated steady currents in DAT (*Fig.3D* and *Fig.3G*, respectively). The voltage-dependence of these currents overlaps with that of DAT-mediated APP^+^ uptake (*Fig. 3J*). In SERT, APP^+^ did not induce sufficiently large steady state currents to determine any current-voltage relationship (*cf. Fig. 3F*). However, serotonin induced robust steady state currents (*Fig. 3I*), the amplitude of which was reduced by ~50% reduction at + 30 mV (diamond symbols, Fig. *3L*). It was not possible to do this comparison in NET (*Fig. 3K*), because neither norepinephrine nor APP^+^ (*cf. Fig. 3E* and *3H*) elicited steady state currents.

**Fig. 3.**
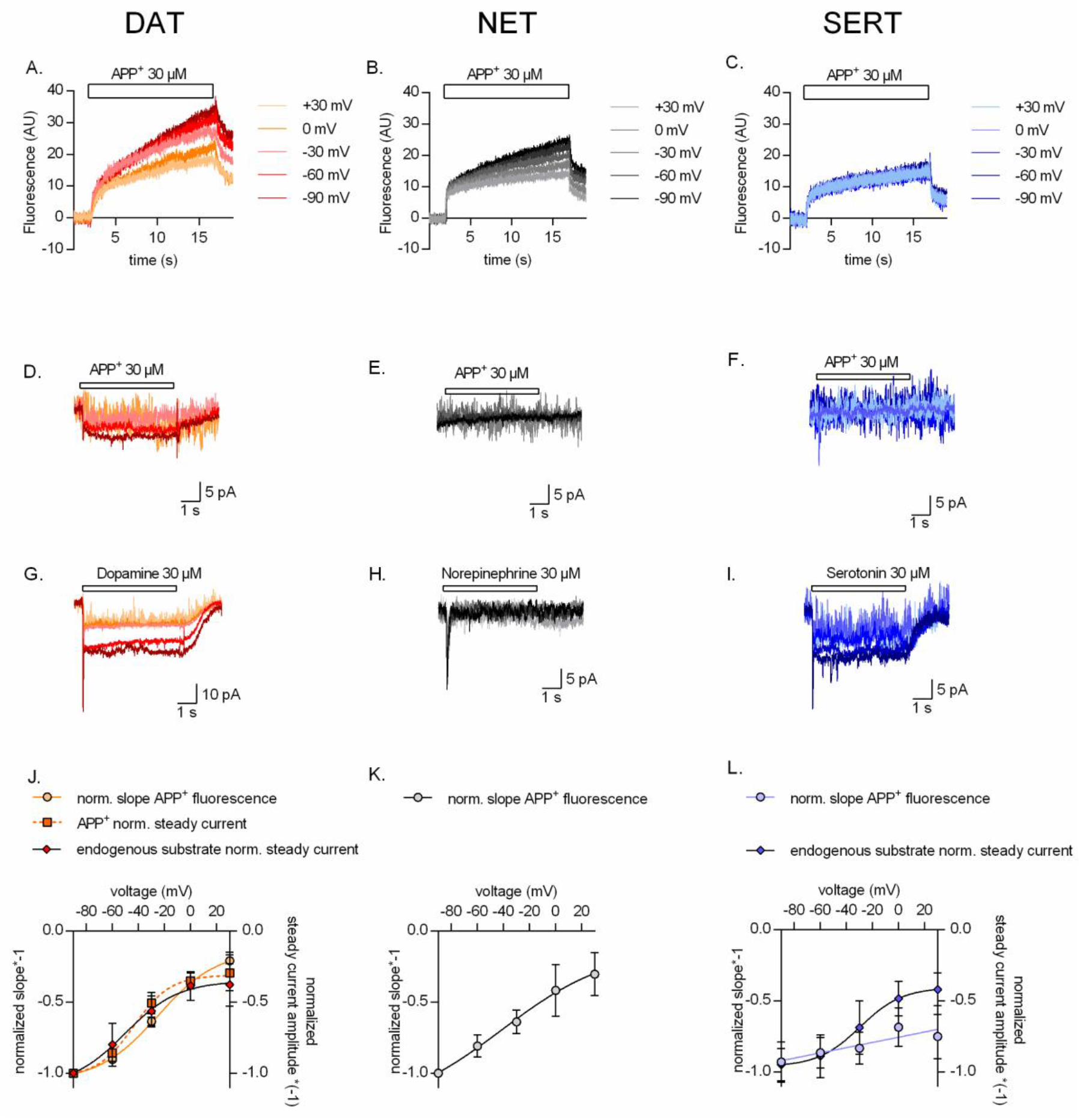
Representative single cell traces of APP^+^-induced fluorescence (*A-C*) and concomitant currents (*D-F*) under different voltages in DAT, NET and SERT, respectively. APP^+^ was applied at concentrations of 30 μM. Representative single cell traces of currents induced by dopamine (30 μM) on DAT (*G*), norepinephrine (30 μM) on NET (*H*) and serotonin (30 μM) on SERT (*I*) under different voltages. (*J*) Normalized voltage dependency of APP^+^-induced fluorescence (circle symbols, n = 6), APP^+^-induced steady state currents (square symbols, n = 6), and dopamine-induced steady state currents (diamond symbols, n = 6) in DAT-expressing HEK293 cells. (*K*) Normalized voltage dependency of APP^+^-induced fluorescence (circle symbols, n = 6) in HEK293 cells expressing NET. Neither norepinephrine nor APP^+^ induced any steady state currents in NET. (*L*) Normalized voltage-dependency of APP^+^-induced fluorescence (circle symbols, n = 6) and serotonin-induced steady state currents (diamond symbols, n = 11) in SERT-expressing HEK293 cells. APP^+^ did not induce any steady state currents in SERT. All experiments were performed under physiological ionic conditions. All fluorescence and current amplitudes were normalized to those obtained at −90 mV (which was set to 1) and the data points were fitted with to the Boltzmann equation (except APP^+^ uptake by SERT, which was fit to a line). All datasets are represented as means ± S.D. AU – arbitrary units, norm. – normalized. We note that the sigmoidal Boltzmann and the line function are both arbitrary fits to the data. Neither one of them, is suitable to model the processes, which underlie the depicted voltage dependence. The decision to use one or the other was based on the fidelity of the resulting fit.

### Impact of intracellular cations on APP^+^ uptake

It is safe to conclude from the observations summarized in *Fig. 3* that NET and DAT differ from SERT in their susceptibility to voltage: transport of APP^+^ by NET and DAT is voltage-dependent; in contrast, influx of APP^+^ mediated by SERT is essentially independent of voltage. Previous studies showed that K_in_^+^ was antiported by SERT, but not by NET or DAT (**Rudnick & Nelson, 1978**; **Gu et al., 1996; Erreger et al., 2008**). Hence, we surmised that differences in the interaction of intracellular K^+^ (K_in_^+^) with DAT, NET and SERT may account for the divergent uptake-voltage relation of DAT or NET and of SERT. Accordingly, we varied the intracellular ionic conditions via the patch pipette and compared the rise in APP^+^ fluorescence over time (represented as AU/sec) in cells expressing DAT, NET and SERT. The uptake-voltage relationship of DAT (*Fig. 4A*), NET (*Fig. 4B*) and SERT (*Fig. 4C*) was similar when measured in the presence of high internal potassium (K_in_^+^ = ~163 mM, circle symbols) or in the absence of potassium (K_in_^+^ = 0 mM, square symbols). In addition and unsurprisingly, raising internal sodium to 163 mM reduced uptake by all three transporters (diamond symbols, *Fig. 4D-F*), because high Na_in_^+^ precludes substrate dissociation on the intracellular side, thus hampering progression of the physiological transport cycle.

**Fig. 4.**
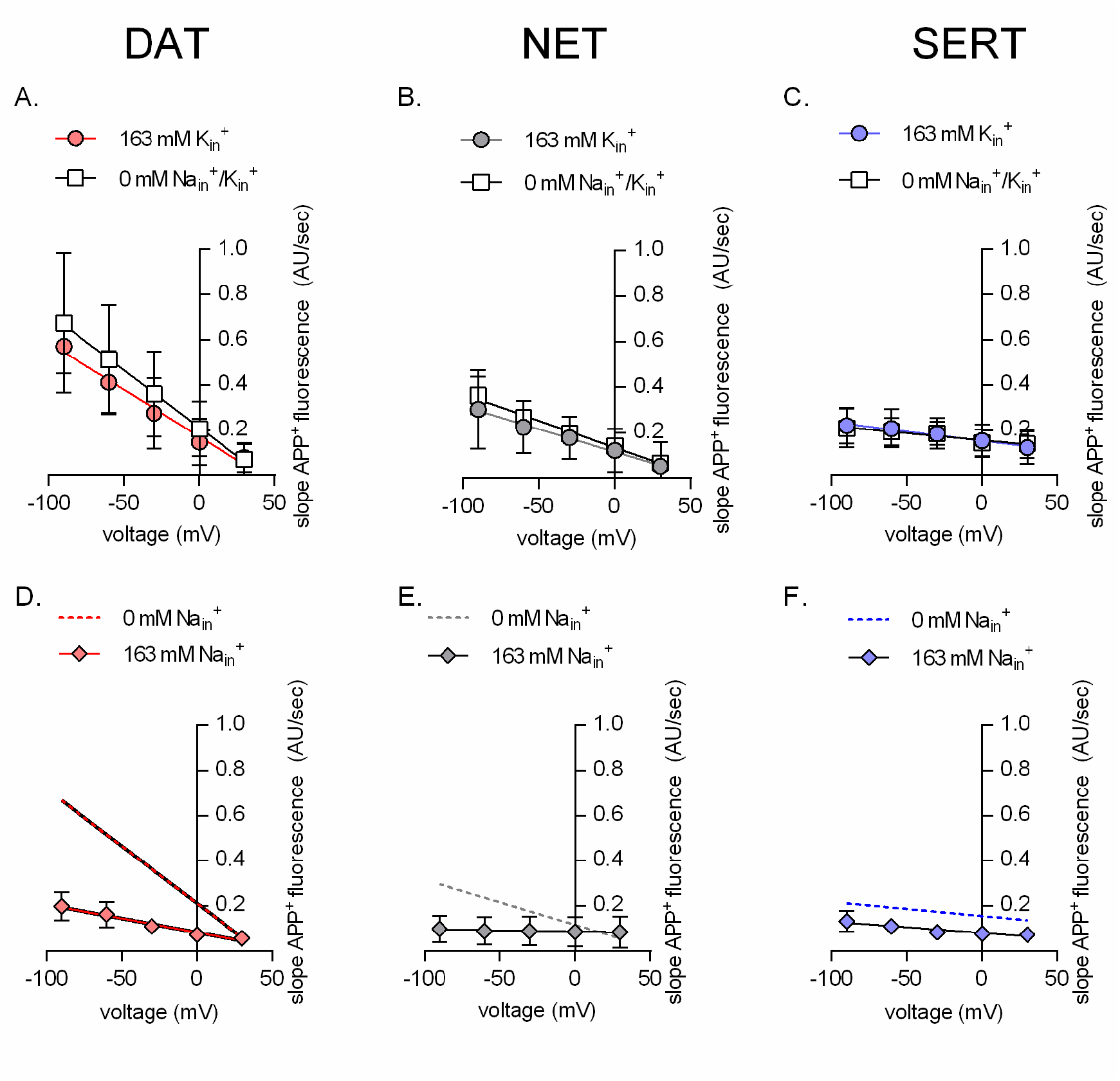
Rate of APP^+^ associated fluorescent uptake (total rise in absolute fluorescence/unit time – AU/sec) measured in DAT (*A* and *D*), NET (*B* and *E*) and SERT (*C* and *F*) under different intracellular conditions and different voltages. Pipette solutions include the physiological intracellular ionic conditions (163 mM K_in_^+^, circle symbols), an intracellular condition devoid of Na_in_^+^ and K_in_^+^ (0 mM Na_in_^+^ /K_in_^+^, square symbols) or an intracellular environment of high Na^+^ (163 mM Na_in_^+^, diamond symbols). All data points were fit to the line equation. The dashed lines in *D*-*F* are the same as those obtained by fitting data points obtained from 163 mM K_in_^+^ from *A*-*C* for the respective transporters. The slope of the lines in the absence and presence of 163 mM K_in_^+^ in *A* (DAT), *B* (NET) and *C* (SERT) were not significantly different (F-test). The p-values were 0.42, 0.65 and 0.63, respectively. This suggest that intracellular K_in_^+^ does not affect the voltage dependence of APP^+^ uptake of either of the three monoamine transporters. The slopes of the lines, in the absence and presence of 163 mM Na_in_^+^, in *D* (DAT) (p = 0.0135) and *E* (NET) (p < 0.0001) were different (F-test). In *F* (SERT), the slope was not significantly different (p = 0.66), while the y intercept was (p < 0.0001). This analysis is consistent with the idea that intracellular Na^+^ hampers APP^+^ through all three transporters. All datasets are represented as means +S.D.; the number of experiments in each individual dataset was 6; AU – arbitrary units.

### Effect of intracellular cations on currents induced by endogenous substrates

Substrate-induced transporter-mediated currents allow for dissecting partial reactions of the transport cycle (**Erreger et al., 2008; Hasenhuetl et al., 2016**). However, APP^+^ elicited only small currents through SERT and NET. Hence, we relied on endogenous substrates to further analyze the effect of intracellular cations on transporter function. We recorded currents through DAT (*Fig. 5A, D, G*), NET (*Fig. 5B, E, H*) and SERT (*Fig. 5C, F, I*), which were elicited by the cognate substrates, in the presence of 163 mM K_in_^+^ (*Fig. 5A-C*), 0 mM K_in_^+^ /Na_in_^+^ (*Fig. 5D-F*) and 163 mM Na_in_^+^ (*Fig. 5G-I*) at different voltages. In the presence of high intracellular potassium, challenge with cognate neurotransmitters elicited an initial peak current through all three transporters (representative original traces in *Fig. 5A-C*). This current reflects translocation of positively charged substrate and Na^+^ across the plasma membrane, which is hindered at positive potentials. This is reflected in the linear peak-current voltage (IV) relationships (circle symbols in *Fig. 5J-L*). Upon elimination of K_in_^+^ (representative original traces in *Fig. 5D-E*), this IV relation became considerably steeper in DAT and SERT and, to a smaller extent, in NET (diamond symbol in *Fig. 5J-L*). In addition and as previously reported (**Erreger et al., 2008; Schicker et al., 2012**), the absence of K_in_^+^ eliminated steady state currents through SERT (*cf. Fig. 5C vs. 5F*), but not through DAT (*cf. Fig. 5A vs. 5D*).

**Fig. 5.**
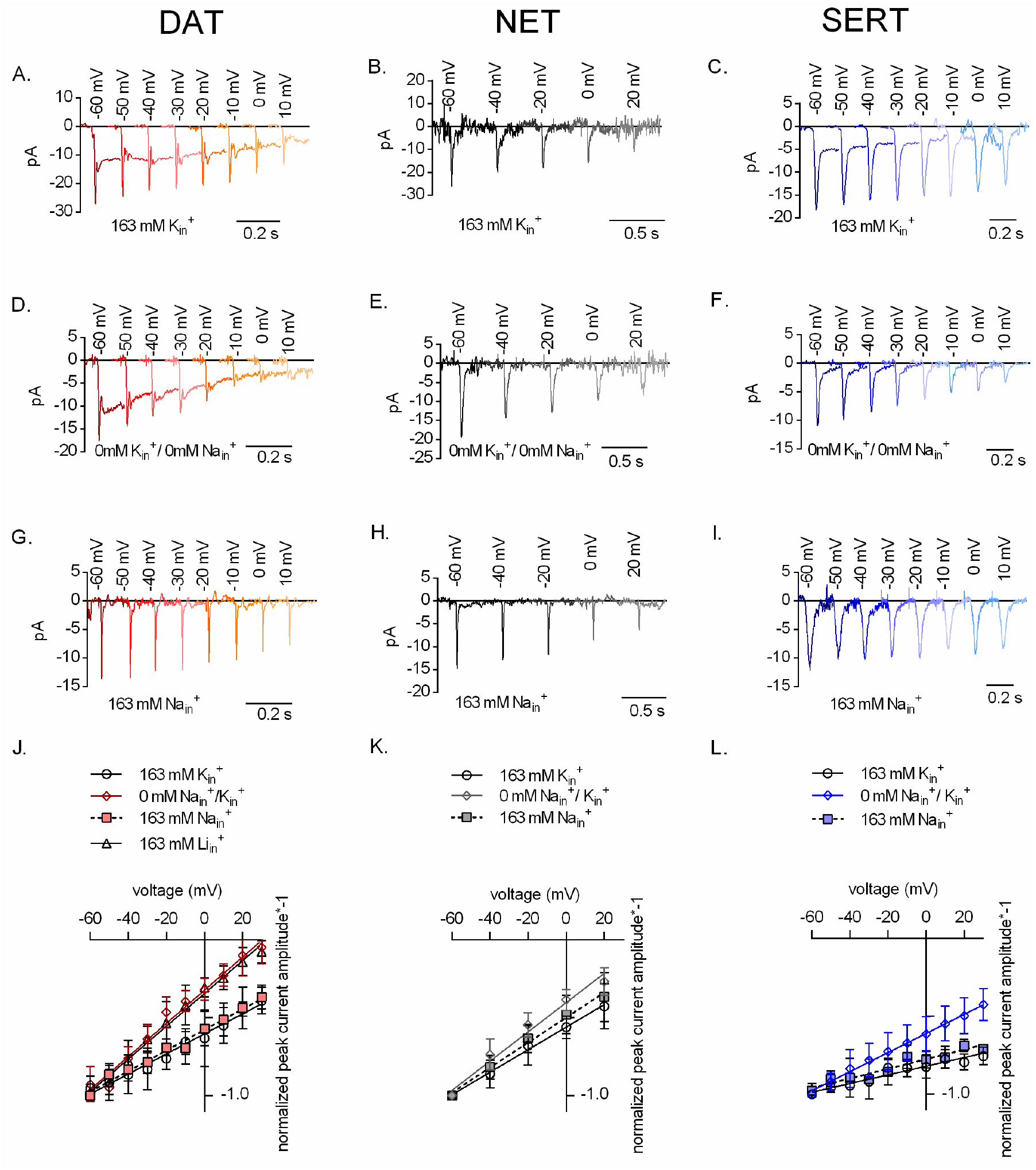
Representative single cell traces of current profiles elicited by 30 μM dopamine in DAT (*A, D* and *G*), 30 μM norepinephrine in NET (*B, E* and *H*) and 30 μM serotonin in SERT (*C, F* and I) under different intracellular conditions and different voltages. Peak current-voltage relationships of DAT (*J*), NET (*K*) and SERT (*L*) under different pipette solutions that include the physiological intracellular ionic conditions (163 mM K_in_^+^, circle symbols), an intracellular condition devoid of Na_in_^+^ and K_in_^+^ (0 mM Na_in_^+^ /K_in_^+^, diamond symbols) or an intracellular environment of high Na^+^ (163 mM Na_in_^+^, square symbols). In the case of DAT (*J*), we also included data obtained with high intracellular Li^+^ (163 mM Li_in_^+^, triangle symbols). Peak current amplitudes were normalized to those obtained at −60 mV (which was set to 1) and the individual datasets were fit with to the line equation. The slope of the voltage dependence in the presence and absence of 163 mM K_in_^+^ was significantly different for all three transporters (DAT (p < 0.0001), NET (p = 0.019) and SERT (p <0.0001); F-test). All data points are represented as means ± S.D.

Because the absence of K_in_^+^ affected the slope of the IV-relation of the peak current, we surmised that potassium bound from the intracellular side not only to SERT but also possibly to DAT and NET. We explored this conjecture by determining the IV relation of peak currents through DAT in the presence of lithium (Li_in_^+^ = 163 mM) instead of K_in_^+^. Li^+^ is an inert cation, because it does not support substrate translocation by *SLC6* transporters and it does not bind to DAT from the intracellular side. The IV relation of peak currents through DAT were similar in the presence of 163 mM L_in_^+^ to those recorded in the absence of K_in_^+^ (triangle symbol *in Fig. 5J*). These observations clearly indicate that K_in_^+^ binds to DAT and rule out an alternative explanation, i.e. that the effect can be accounted for water and monovalent cations briefly occupying a newly available space in the inner vestibule. The IV relations of peak currents are similar in the presence of 163 mM K_in_^+^ (*Fig. 5A-C*) and of 163 mM Na_in_^+^ (*Fig. 5G-I*) (*cf*. circle and square symbols in *Fig. 5J-L*). This is consistent with the idea that Na_in_^+^ and K_in_^+^ bind to overlapping sites in SERT and DAT.

### Effect of K_in_^+^ on uncoupled conductance and catalytic rate of monoamine transporters

Because internal potassium did not affect DAT-mediated uptake (*Fig. 4A*), we examined the role of K_in_^+^ in DAT by determining its effect on transport-associated currents. The presence (square symbol in *Fig. 6A*) and absence of K_in_^+^ (circle symbol in *Fig. 6A*) did not change the voltage dependence of the steady state current. However, the amplitudes of the steady state currents through DAT were smaller in the absence of K_in_^+^. This finding suggests that DAT-mediated currents are not strictly coupled kinetically, i.e., a current component exists, which is uncoupled from the transport cycle. Regardless of the intracellular ion composition, steady state currents were not observed, if NET was challenged with saturating concentrations of norepinephrine (*Fig. 6B*). Currents through SERT (amplitudes of which are represented as diamond symbol in *Fig. 3L* and square symbol in *Fig. 6C*) are completely uncoupled from the catalytic transporter cycle; in spite of its electroneutral stoichiometry, SERT mediates an inwardly directed current, which is eliminated by removal of K_in_^+^ (circle symbol in *Fig. 6C*). These observations are in line with previously published work (**Schicker et al., 2012; Hasenhuetl et al., 2016**). We further confirmed the contribution of K_in_^+^ in the catalytic cycle of all three monoamine transporters by employing a “two-pulse” peak current recovery protocol (**Hasenhuetl et al., 2016**). This protocol relies on the application of the first pulse (reference) of substrate followed by a variable washout interval and the application of a second pulse (test) of the neurotransmitter (representative single cell traces shown in *Fig. 6D*). The first application of substrate results in recruitment of transporters into the transport cycle. Their availability for the second pulse of substrate depends on their completing the transport cycle, i.e. on their releasing the substrate on the intracellular side and their subsequent return to the outward facing conformation. If the intervening washout interval is prolonged, a larger fraction of transporters becomes available for the second substrate pulse and this can be gauged from the progressively larger peak amplitudes as a function of time. Thus, the time course of this recovery provides estimates of the catalytic rate of the transporters. As shown in *Fig. 6E and 6F*, the catalytic rates of DAT and NET in the presence or absence of K_in_^+^ is very similar. In fact, K_in_^+^ fails to render the peak current recovery by DAT voltage-independent (see *supplementary Fig. 1*). In contrast, SERT shows a ~2 fold deceleration in the catalytic rate in the absence of K_in_^+^ when compared to its recovery rate in the presence of high K_in_^+^ (*Fig. 6G*). These observations are in stark contrast to the indistinguishable uptake by SERT of APP^+^ observed in the presence or absence of K_in_^+^ (*Fig. 4C*). This discrepancy can be accounted for by APP^+^ being a poor substrate, an explanation, which is supported by our observations that APP^+^ did not induce any detectable steady state currents in SERT (*Fig. 2F* and *3F*). All three monoamine transporters can also operate in the exchange mode, which is the basis for the actions of amphetamines (Sitte and Freissmuth, 2015). As a control, we examined the recovery in the presence of high Na_in_^+^, which precludes cycling in the forward transport mode and thus forces the transporters into exchange: as predicted, high Na_in_^+^ accelerated the recovery of all three transporters (square symbols in *Fig. 6E-G*).

**Fig. 6.**
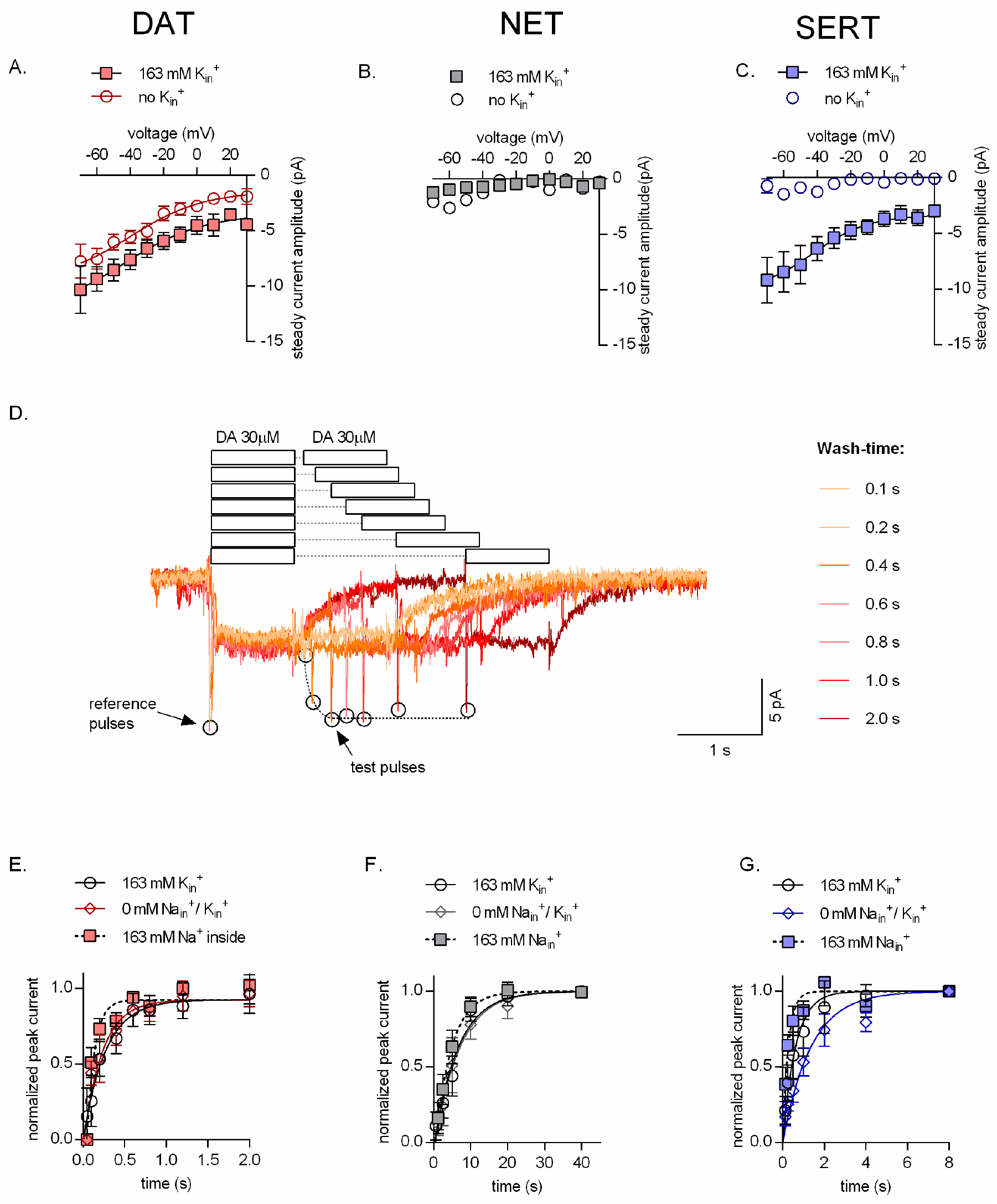
(*A*, *B, C*) Amplitudes of steady state currents elicited by 30 μM dopamine on DAT and 30 μM norepinephrine on NET and 30μM serotonin on SERT, respectively as a function of voltage under physiological intracellular conditions (square symbols) and an intracellular environment devoid of Na^+^ and K^+^ (0 mM Na_in_^+^ /K_in_^+^, circle symbols). NET did not elicit any steady state currents on exposure to norepinephrine. The data points in *A*, *B* and *C* are represented as means ± S.E.M. The lines indicate fits of the Boltzmann equation to the data points. For DAT (*A*), we compared the amplitude of the current in the presence and absence of 163 mM K_in_^+^ at −20 mV. The difference in amplitude at this potential was significant (p= 0.0334); Mann Whitney test (n= 9; each). (*D*) Representative single cell trace of the two pulse protocol: 30 μM dopamine was applied to a DAT-expressing cell to generate a reference pulse followed by variable wash times and repeated pulse of 30 μM dopamine again (test pulse). The peak current amplitudes (which represent available binding sites on completion of the transport cycle) are plotted as a function of time to determine transporter catalytic rate. Catalytic rates determined for DAT (*E*), NET (*F*) and SERT (*G*) under intracellular conditions that are physiological (163 mM K_in_^+^, circle symbols), devoid of intracellular Na^+^ or K^+^ (0 mM Na_in_^+^ /K_in_^+^, diamond symbols) or contain high intracellular Na^+^ (163 mM Na_in_^+^, square symbols). Peak current amplitudes obtained at each test pulse were normalized to that of the reference peak (which was set to 1) and fitted with a mono-exponential function. The catalytic rates obtained were as follows: DAT (163 mM K_in_^+^ – 3.74 ± 0.76 s^−1^; 163 mM Na_in_^+^ – 10.68 ± 2.14 s^−1^; 0 mM K_in_^+^ /Na_in_^+^ – 4.45 ± 0.93 s^−1^), NET (163 mM K_in_^+^ – 0.15 ± 0.013 s^−1^; 163 mM Na_in_^+^ – 0.22 ± 0.016 s^−1^; 0 mM K_in_^+^ /Na_in_^+^ – 0.15 ± 0.014 s^−1^) and SERT (163 mM K_in_^+^ – 1.61 ± 0.12 s^−1^; 163 mM Na_in_^+^ – 2.57 ± 0.28 s^−1^; 0 mM K_in_^+^ /Na_in_^+^ – 0.74± 0.058 s^−1^). The data in *E*, *F* and *G* are represented as means ±_S.D (n = 5 for each condition).

### A kinetic model for the transport cycle of monoamine transporters

The data, represented in *Fig. 6*, can be explained by a model, which posits that all monoamine transporters can bind K_in_^+^, but that the bound K_in_^+^ is released on the intracellular side prior to the return step by DAT and NET. In contrast, K_in_^+^ is released on the extracellular side after being antiported by SERT. We tested the plausibility of this hypothesis by resorting to kinetic modeling. As a starting point for modeling DAT and NET, we used the previously proposed kinetic model for DAT by Erreger and coworkers (**Erreger et al., 2008,**shaded in grey in *Fig. 7A*), which is nested within our proposed model. For NET, we posited a much slower dissociation-rate for the substrate (indicated as green text) to account for the small substrate turnover rate and the absence of the steady current component (cf. *Fig.2H* and *Fig.6E*). The model was expanded to account for transient binding of K_in_^+^ to DAT and NET. This was achieved by subdividing this event into two consecutive reactions: in the first reaction (when viewed in the clockwise direction), DAT/NET adopts an inward-facing conformation on the trajectory to the occluded state (ToccClK), to which K_in_^+^ can still bind, but with reduced affinity. In the second reaction, DAT/NET fully occludes after shedding off K_in_^+^ (i.e., ToccCl) and rearranges to adopt the outward-facing conformation.

**Fig. 7.**
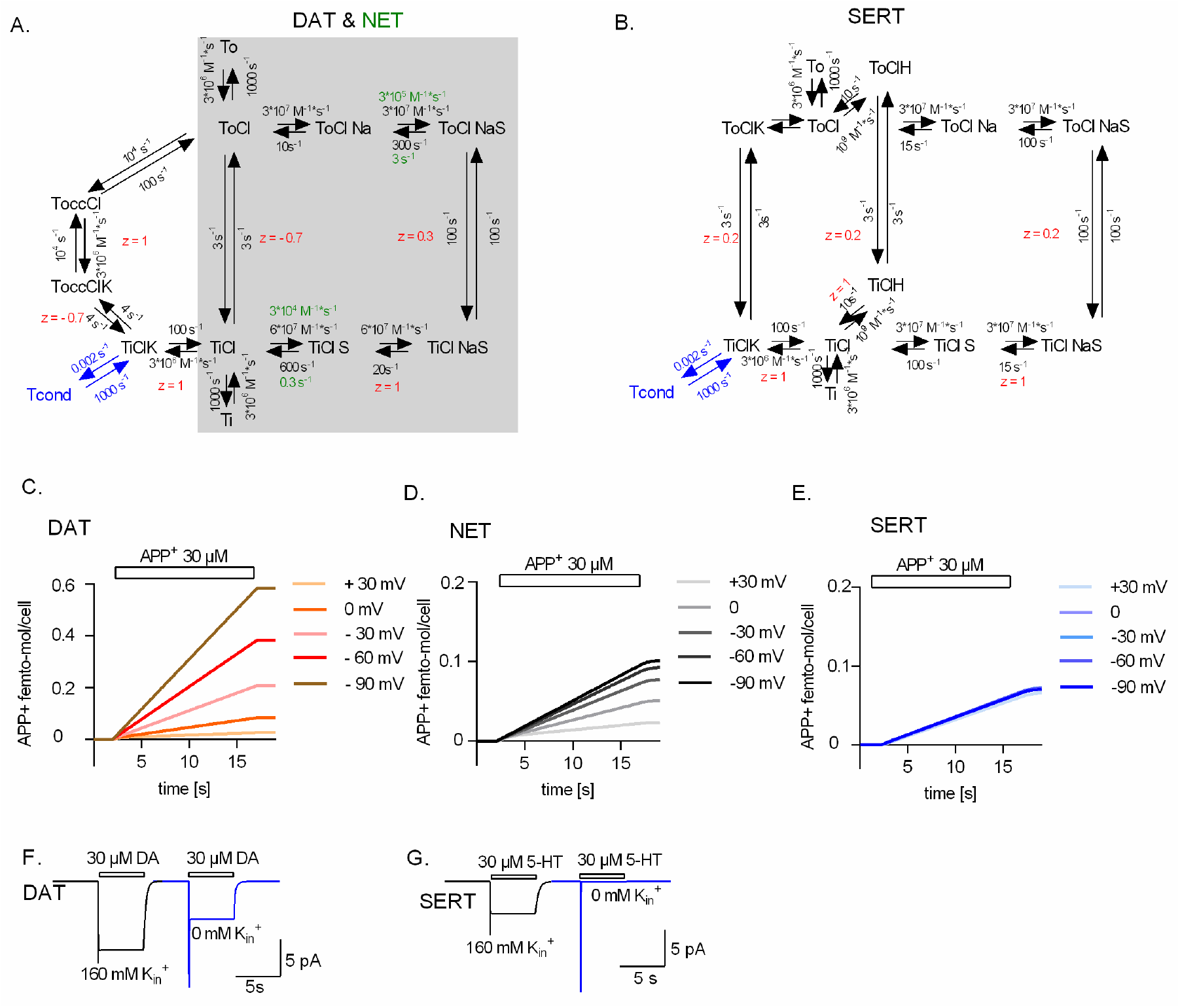
(*A*) Reaction schema of DAT and NET. Shaded in grey is the original schema proposed for DAT by **Erreger et al., 2008**. The original schema is nested in the refined model. We assumed that DAT and NET not only share the same schema but also most parameters. However, to account for the smaller turn-over rate of NET and the absence (or lack of detection) of the steady current component on challenge with norepinephrine, we posited slower substrate binding kinetics for NET (rate constants indicated in green) (**B**) Reaction schema of SERT. Simulated APP^+^ uptake through DAT (*C*), NET (*D*) and SERT (*E*). k_on_ and k_off_ for APP^+^ were set to 9*10^5^ s^−1^ *M^−1^ and 30 s^−1^, 3*10^5^ s^−1^ *M^−1^ and 1 s^−1^ and 1*10^5^ s^−1^ *M^−1^ and 1 s^−1^ for DAT, NET and SERT, respectively. Simulated substrate induced currents for DAT (*F*) and SERT (*G*) in the presence (black trace) and absence (blue trace) of 163 mM K_in_^+^. The current in the presence of 163 mM K_in_^+^ is the sum of the coupled and uncoupled current. The current in the absence of K_in_^+^ is the coupled current in isolation. In simulations (*F*) and (*G*), we assumed a membrane voltage of – 60 mV.

In the model of SERT (originally proposed in **Schicker et al., 2012**; *Fig. 7B*), we assumed that K_in_^+^ was antiported on return of the substrate-free transporter to the outward facing conformation. In both models (i.e., DAT/NET and SERT), we accounted for the uncoupled current component observed in the presence of K_in_^+^ by adding a conducting state, which was in equilibrium with the K_in_^+^-bound inward-facing conformation (Tcond, indicated in blue). Notably, voltage independent substrate transport by SERT in the absence of K_in_^+^ (*Fig. 4C*) was accounted in the model by the ability of SERT to alternatively antiport a proton (**Keyes and Rudnick, 1982; Hasenhuetl et al. 2016**). However, varying intracellular protons did not alter the voltage dependence of the peak current amplitude in DAT (*Supplementary Fig.2*). This finding is consistent with the concept that only SERT can utilize protons in the return step.

We interrogated the kinetic models to generate synthetic data for APP^+^ transport by DAT (*Fig. 7C*), NET (*Fig. 7D*) and SERT (*Fig. 7E*) at different membrane voltages. The synthetic data generated through the respective kinetic models could faithfully reproduce our experimental findings: i) only APP^+^ uptake by SERT was voltage-independent (cf. *Fig.7C-7E* and *Fig. 3A-3C*); ii) the removal of K_in_^+^ abrogated the steady-state current only in SERT but not in DAT (cf. *Fig.7F/7G* and *Fig. 6A/6C*); iii) the removal of K_in_^+^ did not slow down the return of DAT and NET from the inward-to the outward-facing conformation, while it reduced this rate in SERT by two-fold (cf. *Fig. 6E – 6G* to *Supplementary Fig.3C*). For other simulated datasets, please refer to *Supplementary Fig.3*. This indicates that the underlying assumptions are valid and allow for a reasonable approximation, which has explanatory power: the differences in handling of K_in_^+^ incorporated into the model are necessary and sufficient to account for the differences in the forward transport mode of DAT, NET and SERT.

## Discussion

Fluorometric analyses of the SERT, DAT and NET transport cycle provide mechanistic and functional insights into the *modus operandi* of these transporters by combining advantages of assays that employ radiolabeled ligands (used to determine global transporter turnover rates) with those that involve electrophysiology (which provide voltage control and unmatched temporal resolution) (**Schwartz et al., 2006**). Fluorescent substrates of monoamine transporters have previously been used to monitor transporter-mediated uptake in real time (**Schwartz et al., 2003; Mason et al., 2005; Schwartz et al., 2005; Oz et al., 2010; Solis Jr. et al., 2012; Karpowicz Jr et al., 2013; Wilson et al., 2014; Zwartsen et al., 2017; Cao et al., 2020**). One such fluorescent substrate is APP^+^, a fluorescent analog of MPP^+^, a neurotoxin which targets monoaminergic neurons (**Javitch et al., 1985; Scholze et al., 2002**). Like MPP^+^, APP^+^ is also taken up by cells expressing DAT, NET and SERT; its fluorescent properties are well understood (**Solis Jr. et al., 2012; Karpowicz Jr et al., 2013; Wilson et al., 2014**). In the present study, we relied on APP^+^ to explore the transport cycle of DAT, NET or SERT by single cell analysis, which allowed for simultaneously recording cellular uptake by fluorescence and substrate-induced currents. It was also possible to control the concentrations of the relevant ions and the membrane potential with the unprecedented precision of the whole cell patch-clamp configuration and to thereby examine their impact on transport rates. To the best of our knowledge, our experiments are the first to address the following question: why do the structurally similar DAT, NET and SERT differ in transport kinetics and handling of co-substrate binding? It is evident that DAT and NET resemble SERT in most aspects. We show here that all major differences can be accounted for by the distinct handling of K_in_^+^ : (i) in SERT, physiological K_in_^+^ concentrations accelerated the rate of substrate uptake: it was 2-fold faster than in the absence K_in_^+^ (*Fig. 6G*). In contrast, DAT and NET return to the outward-facing state with the same rate regardless of whether K_in_^+^ is present or not (*Fig. 6E* and *6F*). Accordingly, K_in_^+^ did not affect rate of substrate uptake by DAT and NET (*Fig. 4A* and *4B*). (ii) The catalytic rate of SERT was independent of voltage in the presence of physiological ionic gradients (*Fig. 3L* and *Fig. 5L*). This was not the case for DAT and NET (*Fig. 3J* and *3K* and *Fig. 5J* and *5K*, respectively). (iii) In all three transporters, release of Na_in_^+^ from the inward facing conformation is electrogenic. In SERT, this electrogenic Na_in_^+^ dissociation is cancelled out by electrogenic K_in_^+^ binding to the inward-open empty transporter and its subsequent antiport, thereby rendering the cycle completion rate voltage-independent. In DAT and NET, however, the cycle completion rate remained voltage-dependent despite the fact that K_in_^+^ also bound in a voltage-dependent manner. (iv) K_in_^+^ is also relevant to account for the distinct nature of the steady-state current component in SERT and DAT. The steady current component carried by the electroneutral SERT is produced by an uncoupled Na^+^ flux through a channel state that is in equilibrium with the K_in_^+^-bound inward-facing conformation (**Schicker et al., 2012**). This uncoupling explains the existence of differences in the voltage-dependence of SERT-mediated uptake and of steady state currents (*Fig. 3L*). DAT-mediated transport is accompanied by the translocation of coupled net positive charges in each cycle. Thus, DAT-mediated steady-state currents were originally modelled to be strictly coupled to substrate transport (**Erreger et al., 2008**). However, our data suggests that DAT also carries a previously suggested uncoupled current component (**Sonders et al., 1997; Sitte et al., 1998; Carvelli et al., 2004**), which adds to those associated with DAT-mediated ionic transport. This uncoupled current in DAT, just like in SERT, is contingent on the presence of intracellular K_in_^+^. Binding of K_in_^+^ to DAT at the Na2 site was proposed earlier by a study that employed extensive molecular dynamic simulations to understand intracellular Na^+^ dissociation from DAT (**Razavi et al., 2017**). Our results highlight the fact that the voltage dependence of peak amplitudes is identical in the presence of high Na_in_^+^ and high K_in_^+^ (*Fig. 5J*) and thus support this conjecture.

The most parsimonious explanation for all differences between SERT, NET and DAT was to posit that K_in_^+^ is antiported by SERT but not by DAT and NET. All three transporters carry a negative charge through the membrane on return from the substrate-free inward-to the substrate-free outward-facing conformation (presumably a negatively charged amino-acid). In the case of SERT, however, the charge on the transporter is neutralized by the counter-transported K_in_^+^. Because the return step is slow and therefore rate-limiting, it determines the voltage dependence of substrate uptake. The K_in_^+^-binding site in SERT, alternatively, can also accept protons (**Keyes and Rudnick, 1982**). Hence protons – as alternative co-substrate that is antiported – support the return step from the inward-to the outward facing substrate-free conformation (**Hasenhuetl et al., 2016**). The alternative is to postulate, based on recent evidence in LeuT (**Billesbølle et al., 2016**), that antiport of K_in_^+^ is a general feature of all *SLC6* transporters. However, this can be refuted for DAT and NET for the following reasons: the presence or absence of K_in_^+^, did not change their catalytic rates (*Fig.6E* and *6F*). Thus, in the absence of K_in_^+^, the transporters seem to return from the substrate-free inward-to the substrate-free outward-facing conformation. In this case, however, the transporters carry one positive charge less through the membrane. This change in ion translocation must, therefore, translate into a concomitant, substantial change in the voltage dependence of substrate transport, i.e., the voltage dependence of transport ought to be much steeper in the absence than in the presence of K_in_^+^. This was not observed (*Fig.4A* and *4B*). Additionally, H^+^ failed to accelerate the catalytic rate of DAT (data not shown) and the slope of the IV-curve for the peak current remained steep (*Supplementary Fig.2*). These observations indicate that protons (like K_in_^+^) cannot be antiported by DAT.

In monoamine transporters, there is a continuum between full substrates, partial substrates, atypical inhibitors and typical inhibitors (**Hasenhuetl et al., 2019, Bhat et al., 2019**). Interestingly, APP^+^ is a full substrate of DAT: the currents, which were elicited by APP^+^, were indistinguishable to those induced by the cognate substrate dopamine and other full substrates such as D-amphetamine (**Erreger et al., 2008**). In contrast, APP^+^ elicited the peak current but failed to induce the steady current through SERT, which was readily seen in the presence of 5-HT. In oocytes expressing SERT, APP^+^ elicited currents reached only ~20% of the amplitudes of the 5-HT-induced currents (**Solis Jr., et al., 2012**). In SERT expressing HEK293 cells, the cognate substrate elicits currents of a magnitude in the low pA range. Hence, transport-associated currents induced by APP^+^ are expected to be lost in the noise. Taken together these observations, APP^+^ is a poor substrate for SERT: its actions can be rationalized by assuming that it traps transporters in one or several conformational intermediates, which are exited with a rate slower than the return step. Therefore DAT and SERT diverge in their handling of APP^+^ : while APP^+^ had similar K_M_-values for DAT and SERT, the catalytic rate of the transport cycle was equivalent to that of the cognate substrate in DAT, but substantially lower in SERT.

Originally, NET expressed in HEK293 cells was reported to support both, a peak and a steady current, when challenged with substrate (**Galli et al., 1995**). However, in the present study, we only observed the peak current with our superfusion system, which allowed for rapid exchange of solutions: neither APP^+^ nor the cognate substrate norepinephrine elicited a steady current. The absence of the steady current can be attributed to the very slow catalytic rate of NET: it is evident that, by contrast with DAT and SERT, NET returns on a timescale of seconds from the inward-to the outward-facing state: NET dwells in the inward-facing state with a lifetime of τ=~7s (cf. *Fig.6F*), which is >10-20 times longer than the dwell time of DAT and SERT (DAT, τ= ~0.3s; SERT, τ= ~0.6s, *Fig. 6E* & *6G*). Thus, this very slow turnover explains the absence of coupled or uncoupled NET-mediated currents in spite of the proposed electrogenic stoichiometry (**Gu et al., 1996**).

Our analysis provides a unifying concept of substrate transport through all three monoamine transporters, i.e., they are equivalent in all aspects of their transport cycle but one: in SERT, the binding site for K_in_^+^ remains intact upon conversion of the transporter from the inward to the outward facing conformation. In contrast, this binding site is less stable in DAT and NET. The resulting loss in affinity leads to the shedding of K_in_^+^ prior to the return step. The repercussions of this subtle difference are profound: SERT and DAT/NET differ (i) in their voltage-dependence of substrate uptake, (ii) in the nature of the substrate-induced current and (iii) in the energy sources tapped for concentrative substrate transport. If DAT and NET do not antiport K_in_^+^, its concentrative power must be independent of the existing K^+^ gradient across the plasma membrane. On the other hand, if the stoichiometry of DAT and NET are electrogenic, a change in membrane voltage is predicted to increase or decrease substrate uptake at steady state depending on the direction of the voltage change. In this context, it is important to note that the experiments conducted in the present study all report on substrate transport at pre-steady state. Additional insights on whether or not DAT/NET can antiport K_in_^+^ can come from experiments conducted at the thermodynamic equilibrium. Such experiments need to be performed by, for instance, employing a vesicular membrane preparation that contains reconstituted DAT or NET. Such preparations allow for the control of the inner and outer ion composition while preventing the substrate to escape from the vesicular confinement. However steady-state assessment of transporter mediated substrate uptake is hindered by the fact that all three monoamine transporters can also transport substrate in the absence of K_in_^+^. These observations are difficult to reconcile with the concept of transport by fixed stoichiometry. We, therefore, surmise that DAT, NET and SERT operate with a mixed stoichiometry. Based on our data we conclude that DAT and NET are less likely than SERT to antiport K_in_^+^, because we cannot rule out that they can occasionally carry the K_in_^+^ ion through the membrane. Conversely, SERT antiports K_in_^+^ in the majority of its cycles but may return empty in some instances. We thus believe that the differences between these three transporters with respect to their handling of K_in_^+^ represents a continuum, as opposed to divergence, in ionic coupling and kinetic decision points during substrate transport. The difference between SERT and DAT/NET represent different approaches to an inherent trade-off and may reflect an adaptation to physiological requirements: because of electrogenic binding and subsequent counter-transport of K_in_^+^, SERT operates in the forward transport mode with a constant rate regardless of membrane potential, but it cannot exploit the membrane potential to fuel its concentrative power. In contrast, DAT and NET can harvest the energy of the transmembrane potential to fuel its concentrative power. As a trade-off, the substrate uptake rate of DAT and NET is voltage-dependent and strongly reduced or increased upon membrane depolarization or hyperpolarization, respectively.

## Experimental procedures

### Whole cell patch clamping

Whole cell patch clamp experiments were performed on HEK293 cells stably expressing DAT, NET or SERT. These cells were grown in Dulbecco’s Modified Eagle Media (DMEM) supplemented with 10% heat-inactivated fetal calf serum (FBS), 100 u·100 mL^−1^ penicillin, 100 u·100 mL^−1^ streptomycin and l00 μgmL^−1^ of geneticin/G418 for positive selection of transporter expressing clones. Twenty-four hours prior to patching, the cells were seeded at low density on PDL coated 35 mm plates. Substrate-induced transporter currents were recorded under voltage clamp. Cells were continuously superfused with a physiological external solution that contains 163 mM NaCl, 2.5 mM CaCl_2_, 2 mM MgCl_2_, 20 mM glucose, and 10 mM HEPES (pH adjusted to 7.4 with NaOH). Pipette solution mimicking the internal ionic composition of a cell (referred to as normal internal solution henceforth) contained 133 mM potassium gluconate, 6 mM NaCl, 1 mM CaCl_2_, 0.7 mM MgCl_2_, l0 mM HEPES, 10 mM EGTA (pH adjusted to 7.2 with KOH, final K_in_^+^ concentration – 163 mM). A low Cl^−^ internal solution was made by replacing NaCl, CaCl_2_ and MgCl_2_ in normal internal solution by NaMES, CaMES and Mg-Acetate (MES – methanesulfonate). A high Cl^−^ internal solution was made by replacing potassium gluconate in the normal internal solution with KCl. Na_in_^+^ and/or K_in_^+^-free internal solutions were made by replacing NaCl and/or potassium gluconate respectively in the normal internal solution with equimolar concentrations of NMDG chloride (titrated to pH of either 7.2 using NMDG or with KOH in Na_in_^+^-free 163 mM K_in_^+^ internal solution. An internal solution with high Na_in_^+^ concentration was made by replacing potassium gluconate of the normal internal solution with equimolar concentration of NaCl (pH adjusted to 7.2 with NaOH). A high Li^+^ internal solution was made by replacing potassium gluconate in the normal internal solution with 130 mM of LiCl (pH adjusted to 7.2 with LiOH, final Li_in_^+^ concentration – 163 mM). Internal solution with a pH of 5.6 was prepared with 10 mm 2-(N-morpholino)ethanesulfonic acid buffer, 1 mm CaCl2, 0.7 mm MgCl2, 10 mm EGTA, and 140 mm NMDGCl and was titrated to pH 5.6 with NMDG. Currents elicited by dopamine or APP^+^, a fluorescent substrate of DAT (IDT307, Sigma Aldrich), were measured at room temperature (20-24^°^C) using an Axopatch 200B amplifier and pClamp 10.2 software (MDS Analytical Technologies). Dopamine, norepinephrine, serotonin or APP^+^ was applied using a DAD-12 superfusion system and a 4-tube perfusion manifold (ALA Scientific Instruments), which allowed for rapid solution exchange. Current traces were filtered at 1 kHz and digitized at 10 kHz using a Digidata 1550 (MDS Analytical Technologies). Current amplitudes and accompanying kinetics in response to substrate application were quantified using Clampfit 10.2 software (Molecular Devices). Passive holding currents were subtracted, and the traces were filtered using a 100-Hz digital Gaussian low-pass filter.

### Simultaneous fluorescence-current recordings

Twenty-four hours prior to fluorescence recording, HEK293 cells stably expressing DAT, NET or SERT were seeded at low density on PDL-coated 35 mm glass bottom dishes, which have a cover glass (Cellview Cell Culture Dish, Greiner Bio-One GmbH; Germany). On the day of the experiment, individual cells were visualized and patched using a 100x oil-immersion objective under voltage clamp. APP^+^, a fluorescent molecule that has an excitation range from 420-450 nm, was applied to single cells using a perfusion manifold. APP^+^ uptake into the cell was measured using a LED lamp emitting 440 nm light and a dichroic mirror that reflected the light onto the cells. The emitted fluorescence from the sequestered APP^+^ within the cells was measured using photomultiplier tubes (PMT2102, Thorlabs, United States) mounted on the microscope after it had passed an emission filter. The signal from the PMT was filtered at 3 kHz, digitized at 10 kHz with an Axon Digidata 1550B and pClamp 10.2 software (MDS Analytical Technologies). Current traces were filtered as mentioned above. The signals (i.e. currents and fluorescence) were acquired with separate channels.

### Kinetic modelling and statistics

The kinetic model for the DAT transport cycle is based on previously reported sequential binding models for DAT (**Erreger et al., 2008**) and SERT (**Hasenhuetl et al., 2016**). State occupancies was calculated by numerical integration of the resulting system of differential equations using the Systems Biology Toolbox (**Schmidt and Jirstrand, 2006**) and MAT LAB 2017a software (Mathworks). The voltage dependence of individual partial reactions was modeled assuming a symmetric barrier as k_ij_ = k^0^ ije^−zQijFV/2RT^, where F = 96,485 C·mol^−1^, R = 8.314 JK^−1^ mol^−1^, V is the membrane voltage in volts, and T = 293 K (**Läuger, 1991**). Coupled membrane currents upon application of substrate were calculated as I = −F×NC/N_A_×Σ_zQij_(p_i_k_ij_ −p_j_k_ji_), where z_Qij_ is the net charge transferred during the transition, NC is the number of transporters (4 × 10^6^ /cell), and NA = 6.022e^23^ /mol. The substrate-induced uncoupled current was modelled as a current through a Na^+^-permeable channel with I =P_oγ_NC(V_M_ −V_rev_), where Po corresponds to the occupancy of the channel state, γ is the single-channel conductance of 2.4 pS, V_M_ is the membrane voltage, and Vrev is the reversal potential of Na^+^ (+ 100 mV). The extracellular and intracellular ion concentrations were set to the values used in the respective experiments. To account for the non-instantaneous onset of the substrate in patch-clamp experiments, we modeled the substrate application as an exponential rise with a time constant of 10 ms. Uptake of APP^+^ was modeled as TiClS*k_off_S_in_-TiCl*k_o_nS_in_*S_in_* NC/N_A_, where TiClS and TiCl are the respective state occupancies, k_on_S_in_ and k_off_S_in_ are the association and dissociation rate constants of APP^+^ and S_in_ is the intracellular concentration of APP^+^. Experimental variations are either reported as means ± 95% confidence intervals, means ± SD, or means ± SEM. Some of the data was fit to the Boltzmann equation (Y=Bottom+(Top-Bottom)/(1+exp((V50-X)/Slope)) or the line function (linear regression). However both are arbitrary fits to the data. Neither one of them, is suitable to model the processes, which underlie the depicted voltage dependence. The decision to use one or the other was based on the fidelity of the resulting fit.

## Acknowledgments

We thank Verena Burtscher for discussion and comments on the data.

## Footnotes

This work was supported by the Austrian Science Fund (FWF) grant P 31599 and P 31813 to W.S, W1232 (MolTag) to H.H.S and the Vienna Science and Technology Fund (WWTF) grant CS15-033 to H.H.S and LS17-026 to M.F. The authors declare that they have no conflicts of interest with the contents of this article.

## Supplement

**Supplementary Figure 1.**
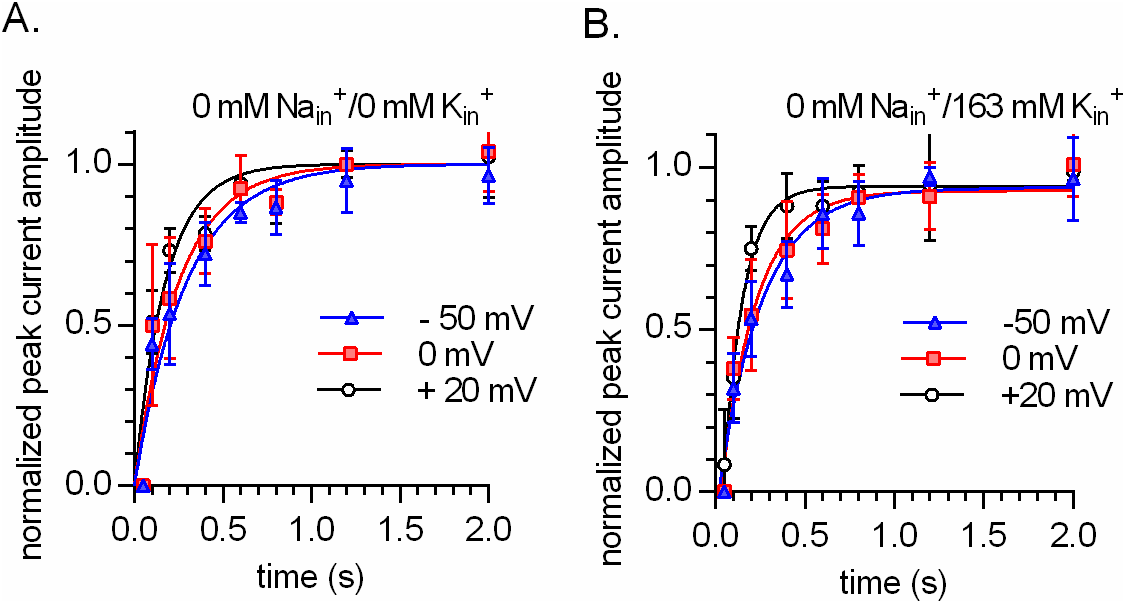
Intracellular K^+^ does not render the time course of peak current recovery of DAT voltage independent. A and B. show the results from two pulse protocols conducted at – 50 mV (triangle), 0 mV (squares) and + 20 mV (circles). The tested conditions were 0 mM Nain K_in_^+^ /0 mM K_in_^+^ in A and 0 mM Nain K_in_^+^ /163 mM K_in_^+^ in B (n = 6; each-error bars indicate SD). The peak current was elicited by the application of 30 μM dopamine. The solid lines in blue, red and black are monoexponential fits to the data. The rates in A. were 3.407 s^−1^ (95% confidence interval: 2.796-4.017 s^−1^), 3.996 s^−1^ (95% confidence interval: 3.056-4.936 s^−1^) and 5.712 s^−1^ (95% confidence interval: 4.470-6.954 s^−1^) for – 50 mV, 0 mV and + 20 mV, respectively. The rates in B. were 3.143 s^−1^ (95% confidence interval: 2.626-3.660 s^−1^), 3.383 s^−1^ (95% confidence interval: 2.734-4.031 s^−1^) and 4.920 s^−1^ (95% confidence interval: 3.800-6.040 s^−1^) for – 50 mV, 0 mV and + 20 mV, respectively.

**Supplementary Figure 2.**
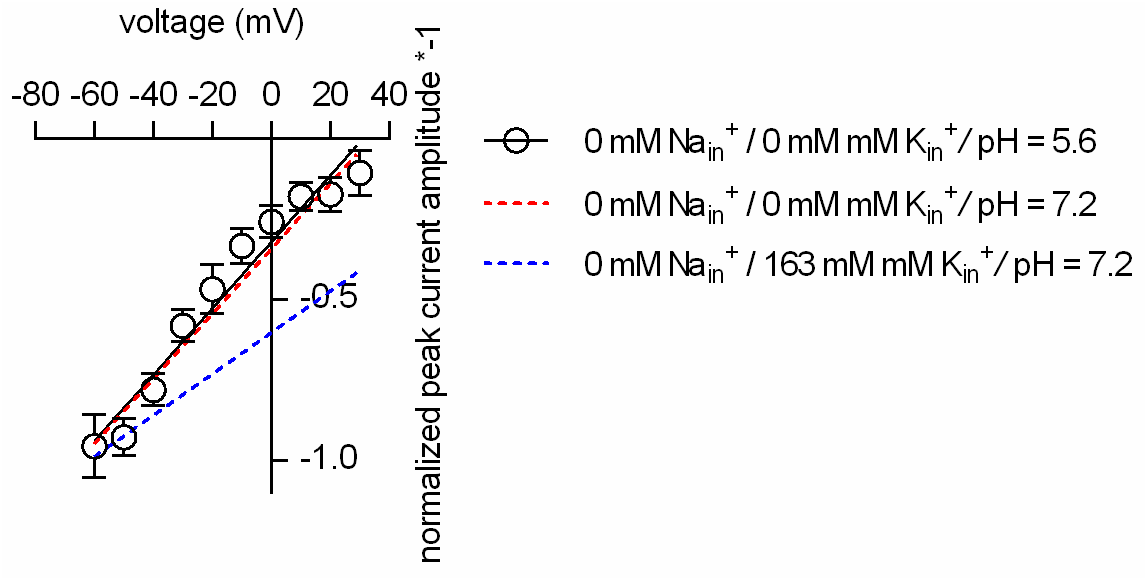
Intracellular protons cannot replace K_in_^+^ in DAT. Plotted is the normalized peak current amplitude as a function of voltage. The peak currents were elicited by rapid application of 30 μM dopamine. The dashed red and blue lines are the fits to the data in *Fig.5J* (main manuscript). At pH 5.6 intracellularly (open circles; n=5; error bars indicate SD) the slope of the voltage dependence remains steep when Na_in_^+^ and K_in_^+^ are absent.

**Supplementary Figure 3.**
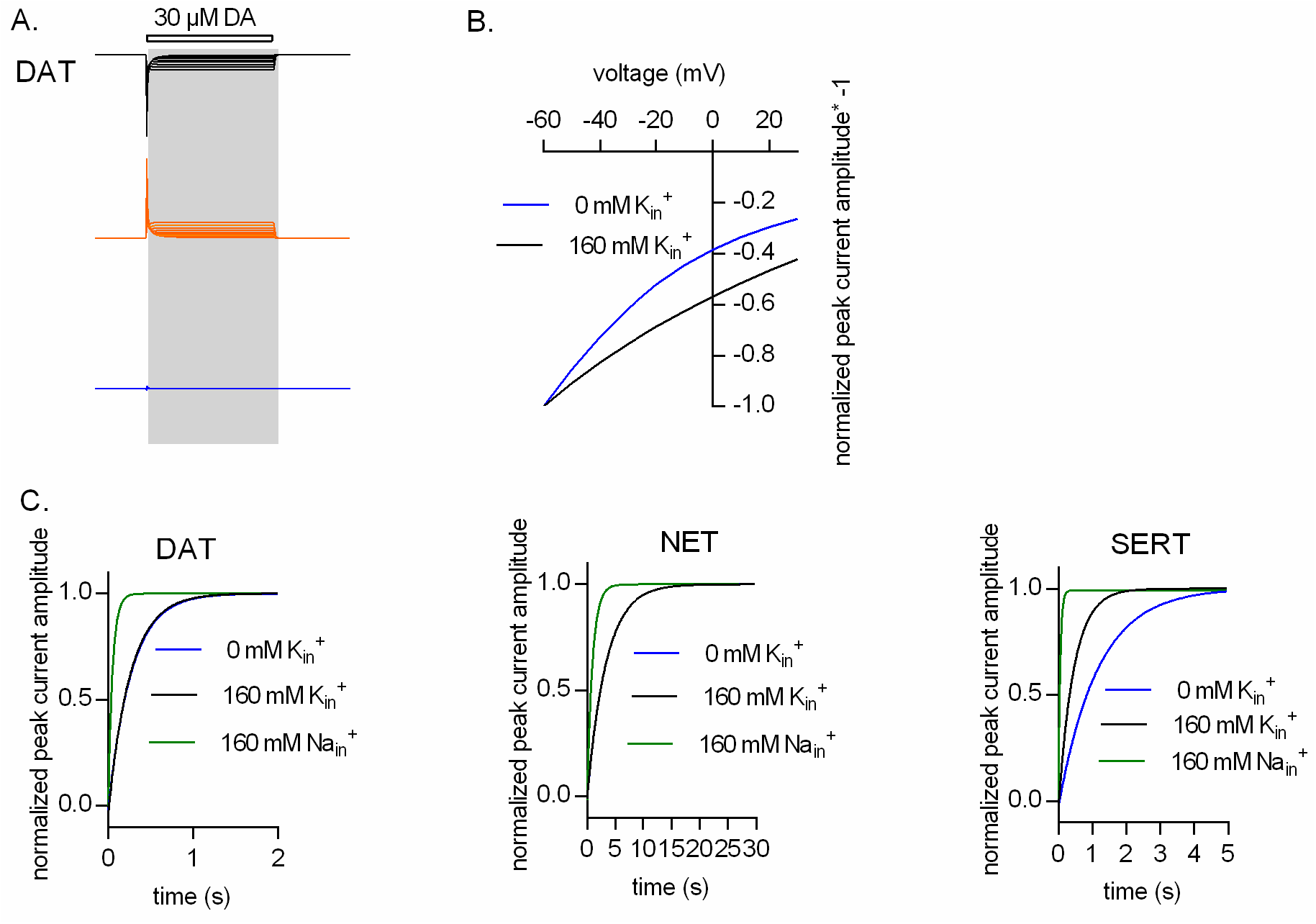
K_in_^+^ binding to inward facing state of DAT affect the voltage dependence of the peak current. A. Our model posits sequential binding and unbinding of Na_in_^+^ and K_in_^+^ to the inward facing conformations and assigns equal valences to these reactions. Thus, the inwardly directed current, produced by outgoing Na_in_^+^ ions dissociating into the cytosol, is canceled out by the outwardly directed current generated by the incoming K_in_^+^ ions. This point is illustrated for DAT., which shows the simulated current components produced by dissociating Na_in_^+^ ions **(black traces, upper panel)**and associating K_in_^+^ ions **(orange traces, middle panel)**over the voltage range from – 60 mV to + 30 mV. Because of the rapid kinetics of ion binding/unbinding, these currents cancel each other out: their sum is zero **(blue trace, bottom panel)**. B. In the absence of **K_m_**^+^ the current, which is generated by Na^+^ ions dissociating into the cytosol is not cancelled out. As a consequence the voltage dependence of the peak current in the absence of K_in_^+^ is steeper than in its presence. C. Simulated peak current recovery in the presence of 160 mM Na_in_^+^ (green line), 160 mM K_in_^+^ (black line) and 160 mM NMDGin^+^ (blue line) for DAT (left panel), NET (middle panel) and SERT (right panel).

